# A Herpes Simplex Virus Type-1-derived influenza vaccine induces balanced adaptive immune responses and protects mice from lethal influenza virus challenge

**DOI:** 10.1101/2021.08.05.455241

**Authors:** Paul J.F. Rider, Harrison Dulin, Ifeanyi K. Uche, Michael C. McGee, Blake Breitenstein, Gene S. Tan, Weishan Huang, Konstantin G. Kousoulas, Rong Hai

## Abstract

Influenza virus is a major respiratory viral pathogen responsible for the deaths of hundreds of thousands worldwide each year. Current vaccines provide protection primarily by inducing strain-specific antibody responses with the requirement of a match between vaccine strains and circulating strains. It has been suggested that anti-influenza T-cell responses, in addition to antibody responses may provide the broadest protection against different flu strains. Therefore, to address this urgent need, it is desirable to develop a vaccine candidate with an ability to induce balanced adaptive immunity including cell mediated immune responses. A live viral vector technology should exhibit safety, immunogenicity, effectiveness in the presence of pre-existing immunity, and the ability to induce mucosal immune responses. Here, we used VC2, an established Herpes Simplex Virus type 1 vaccine vector, to express the influenza HA protein. We show that this virus is capable of generating potent and specific anti-influenza humoral and cell-mediated immune responses. We further show that a single vaccination with the VC2-derived influenza vaccine protects mice from lethal challenge with influenza virus. Our data support the continued development of VC2-derived influenza vaccines for protection of human populations from both seasonal and pandemic strains of influenza. Finally, our results support the potential of VC2-derived vaccines as a platform for the rapid development of vaccines against emerging and established pathogens, particularly respiratory pathogens.

## Introduction

Current approaches to influenza vaccines have limited efficacy and leave human populations susceptible to emerging influenza strains (1, 2). Indeed, since the development of the first influenza vaccine there have been three pandemics responsible for more than 2 million deaths worldwide and seasonal influenza infection is currently responsible for the deaths of 250,000-500,000 individuals per year worldwide (3–5). In order to produce vaccine in time for the influenza A season it is necessary to predict which influenza A strains will be circulating in the future (6). As might be expected this process is imperfect and predicted strains do not always match the circulating strains (6, 7). In addition to mismatch problems seasonal influenza vaccines exhibit 10-60% efficacy and this leaves human populations vulnerable to more severe illness and death from influenza infection (8–10). Finally, current influenza vaccines leave human populations susceptible to emerging influenza strains for which there is no vaccine, as in pandemic scenarios (11). Therefore, a major focus of influenza vaccine research is to generate a vaccine that will protect human populations from circulating as well as potentially emerging pandemic influenza strains, a so-called “universal” influenza vaccine (12, 13).

The incomplete protection provided by current influenza vaccines is due to the unique biology of influenza virus as a segmented RNA virus: 1) RNA virus replication is error-prone and mutations can occur that lead to the emergence of novel influenza A strains, and 2) influenza A viruses can swap segments, which leads to the emergence of novel subtypes of virus (5, 14). Current influenza vaccination strategies induce strain specific neutralizing antibodies directed against the influenza hemagglutinin (HA) protein (15, 16). These antibodies inhibit the receptor binding activity of HA and effectively neutralize virus entry preventing infection (17–22). As mentioned above the virus can: 1) mutate the receptor binding site (drift) and 2) reassort to include a novel HA segment (shift). This inherent ability to shift and drift results in the emergence of influenza strains for which there is no protection in the human population and for which vaccines do not exist (6). As such, current influenza vaccines require constantly updated vaccine components to match viruses which are predicted to be circulating in a given season.

A major focus of influenza vaccine research is the generation of universal vaccines that can protect against all possible strains of influenza. Critical to achieving the goal of a universal vaccine is technology that is capable of eliciting anti-influenza T-cell responses. The goal of such a vaccine is to generate antibodies and/or T-cell responses that are capable of preventing infection by emergent strains of influenza virus regardless of shift or drift mutations (11, 23). Unlike antibody responses, which typically target HA with neutralizing antibodies that are strain specific, T-cells are capable of targeting infected cells using epitopes that are essential, conserved between different strains of a virus and not subject to shift or drift mutations. It has been suggested that development of vaccines capable of inducing anti-influenza T-cells, particularly at mucosal surfaces, is an important goal of efforts to generate more effective influenza virus vaccines (9, 24). More broadly, there is a need for vaccine platforms capable of inducing strong T-cell responses (25).

Live viral-vectored vaccines are seen as desirable for a number of reasons including their rapid adaptability to deliver antigenic molecules from emerging and re-emerging pathogens, their immunogenicity, and their ability to induce both antibody and T-cell responses. There are currently a number of viral-vectored vaccines in use in veterinary medicine, as well as in late stage clinical trials for human infectious diseases including leading candidates for Ebola and SARS CoV-2 vaccines (26). While herpesviruses are currently not in clinical use as vaccine vectors, the first FDA approved oncolytic virotherapy, T-Vec^™^ is a Herpes Simplex Virus Type 1 (27). Herpesviruses exhibit a number of characteristics that inform their usage as viral vectors: 1) large genome allows the insertion of a number of antigenic transgenes, 2) molecular virology is relatively well understood which informs rational attenuation strategies, 3) relative safety, and 4) the ease of genetic manipulation. Importantly, in clinical and pre-clinical studies, herpesvirus vectors, including HSV-1, have been shown to be unaffected by pre-existing immunity (28–32).

For the current study we have used the herpesvirus vaccine candidate VC2 as the vector to generate an influenza virus vaccine. VC2 contains mutations in the HSV-1 envelope proteins gK and UL20 (33), which render the virus unable to enter into neurons via axonal termini (34–36). As neurons are the site of HSV-1 latency, the mutation in VC2 preclude the establishment of a latent infection. As the majority of clinical symptoms from HSV-1 infection occur after reactivation from latency the inability of VC2 to establish latency greatly enhances its safety profile. In multiple studies using a viariety of animal models, VC2 has been shown to be a safe, effective vaccine against genital herpes in murine, guinea pig, and non-human primate models (33, 37–41). Notably, these studies described the development of significant, durable HSV-1 mucosal immunity in mice that protected against ocular HSV-1 challenge (39). These studies also described the development of anti-HSV-1 mucosal immunity in guinea pig and non-human primate models (40, 41).

In this study, we introduced influenza PR8 HA gene into VC2 and established an optimized design to achieve the maximal expression of HA protein. We further characterized its immunogenicity in mice, finding a robust, specific induction of anti-HA antibody and T-cell responses. Importantly, T-cell responses exhibited both effector and memory phenotypes. We report here that vaccination of mice with a single dose of our VC2-derived influenza vaccine protects mice from lethal challenge with influenza virus. Our data supports the utility of VC2 as a live-attenuated vaccine vector, which has the potential to be applied to the development of a universal influenza vaccine as well as vaccines against significant emerging and re-emerging pathogens, such as SARS-CoV-2.

## Methods

### Cells and viruses

Vero (African green monkey kidney) cells were purchased from ATCC and cultured based on instruction provided by ATCC, using Dulbecco modified Eagle medium (DMEM) with 10% fetal bovine serum (FBS). All recombinant HSVs were propagated in Vero cells. Vero cell monolayers at 80% confluence were infected with 0.1 PFU/cell. Virus was harvested at three days post infection by subjecting the cell monolayers to two freeze-thawing cycles. Virus titers were determined using standard plaque assays on RS cells as described previously (34).

### Bacteria and Plasmids

The construction and characterization of a bacterial artificial chromosome (BAC) plasmid containing HSV-1 VC2 genome have been described previously (34).

This BAC plasmid was used to construct VC2-HA. Briefly, the new VC2–HA plasmid were constructed in Escherichia coli SW105 cells, using the two-step bacteriophage lambda Red-mediated recombination system, as described previously (15, 46). The pCAGGS-PR8-HA sequence of was amplified by PCR using primers P1 and P2. To construct PR8HA-Kan^r^, the kanamycin resistance (Kanr) gene adjoining the I-SceI site was amplified by PCR from plasmid pEPkan-S using primers P3 and P4, fused with PR8HA through fusion PCR. The PR8HA-Kanr gene was amplified by PCR using primers P5 and P6 and then cloned into VC2 to replace gC. The kanamycin resistance cassette was cleaved after expression of I-SceI from plasmid pBAD-I-SceI. The inserted PR8HA was verified by capillary DNA sequencing by using primers P7 to P12.

### Growth kinetics of recombinant viruses in Vero cells

Vero cells were infected at a multiplicity of infection (MOI) of 0.1 and incubated at 33°C in Dulbecco modified Eagle medium (DMEM) with 5% fetal bovine serum (FBS). Viral titers in supernatants were determined by plaque assay on Vero cells.

### Western blotting Analysis

Western blotting and indirect immunofluorescence analysis. One well of a six-well dish of 80% confluent Vero cells was infected (MOI of 2) with the indicated recombinant HSV viruses or mock infected with phosphate-buffered saline (PBS) for 1 h at 33°C. At 24 h postinfection (hpi), cells were lysed in 1X protein loading buffer as described previously. The reduced cell lysates were subjected to Western blot analysis by using monoclonal antibody against A/PR8/HA (PY102), and ICP5 HSV-1 antibody (ab6509, Cambridge MA), as well as monoclonal anti-actin (Sigma, St. Louis, MO). The final Western blotting bands were visualized using an enhanced chemiluminescence protein detection system (PerkinElmer Life Sciences, Boston, MA).

### Animals, vaccination, and challenge

The animal experiments were performed with 6-to 8-week-old female C57BL/6J mice (the Jackson Laboratory, 000664). Animals were anesthetized for all procedures by administering isoflurane intro-nasally._Mice were intramuscularly vaccinated with 1X10^6^ PFU of VC2 control vector or VC2-HA, followed by a boost three week later. Mice were euthanized 7-14 days after the boost. Three weeks after immunization, mice were challenged by intranasal infection with the influenza A/PR8 H1N1 virus at 1X10^3^ PFU. Survival and body weight loss were monitored for 14 days post PR8 challenge. All animal experiments were performed in full compliance with the guidelines of the Institutional Animal Care and Use Committee, at University of California, Riverside and Louisiana State University.

### Enzyme linked immunosorbent assay (ELISA)

To assess the levels of virus-specific antibodies present in immunized mice, enzyme-linked immunosorbent assays (ELISAs) were performed on diluted serum samples as described earlier (16). Briefly, serum was obtained from mice right before viral challenge and stored at −80°C. We coated 96-well ELISA plates (Immulon4; Dynex, Chantilly, VA) with 50 μl (10 μg/ml) of the purified influenza A PR8 viruses. After being washed with PBS, coated wells were blocked with PBS containing 1% BSA and then incubated with diluted serum. After 1 h of incubation at room temperature, wells were rinsed with PBS and incubated with a secondary anti-mouse IgG conjugated to peroxidase (Invitrogen, Carlsbad, CA). Rinsed wells were incubated with 100 μl of SigmaFast OPD (ophenylenediamine dihydrochloride) substrate (Sigma-Aldrich) for 30 min and stopped with 50 μl of 3M hydrochloric acid. The plates were then read with a plate reader that measured the optical density at 490 nm (OD405; DTX880 multimode detector; Beckman Coulter).

### Micro-neutralization assay

PR8 virus was diluted to 1000 PFU per 50 μl with PBS-BSA and then incubated with a series of dilutions of RDE-treated sera, for 1 h at 37°C. MDCK cells in a 96-well plate format were then washed with 1X PBS and infected with 100μ l of the virus and MAb mixture for 1 h at 37°C, 5% CO2. Cells were washed once with 1XPBS and replaced with 1X MEM supplemented with TPCKtreated trypsin. At 24 hour post infection, cell were fixed and permeabilized with ice-cold 80% acetone and air dried. Cells were blocked with 5% NF milk–1XPBS for another 30 min at RT. A mouse anti-M2 antibody was used as a primary antibody. After 1 h of incubation at room temperature, cell were washed with 1XPBS and incubated with a secondary anti-mouse IgG conjugated (Invitrogen, Carlsbad, CA). Similar to earlier ELISA procedure, the plates were incubated with 100 μl of SigmaFast OPD (ophenylenediamine dihydrochloride) substrate (Sigma-Aldrich) for 30 min and stopped with 50 μl of 3M hydrochloric acid. They will be read with a plate reader that measured the optical density at 490 nm (OD405; DTX880 multimode detector; Beckman Coulter).

### Isolation of Splenocytes

Spleens harvested from euthanized mice were pressed through a 70-μm filter to obtain a single-cell suspension, which was centrifuged (600 rcf for 5 min at 4°C). Recovered cells were further depleted of red blood cells by resuspending cells with 5-ml of AKC lysis buffer (Lonza, Walkersville, MD). Following 5 min incubation at room temperature, complete

RPMI medium (RPMI-1640 supplemented with 2 mM L-GlutaMax [Hyclone], 1X MEM nonessential amino acids [Corning], 100U/ml penicillin-streptomycin solution [Hyclone], 1 mM sodium pyruvate [Corning], 0.01M HEPES solution, and 10% fetal bovine serum) was added to neutralize the ACK lysis buffer and the remaining cells were recovered by centrifugation and further resuspended with 5ml of complete RPMI medium.

### T cell Stimulation

To determine antigen-specific T cell expansion in the vaccinated animals, splenocytes were stimulated with HSV gB peptide (1 μg/ml) or PR8 HA protein (5 μg/ml) in vitro for 7 days. To detect antigen-specific T cell cytotoxicity in the vaccinated animals, splenocytes were stimulated with HSV gB peptide (1 μg/ml) or heat-inactivated PR8 strain influenza virus particles (2×10^8^ particles/ml), while cell stimulation cocktail (Tonbo Biosciences) or medium alone conditions were used as positive and negative controls respectively. Brefeldin A (5 μg/ml, Sigma) and Monensin (2 μM, Sigma) were added 1 hr post stimulation, following incubation at 37°C for 5 h.

### Flow Cytometry Analysis

The following antibodies were used for flow cytometry: allophycocyanin (APC)-CD4 (GK1.5), APC-Cy7-8a (53-6.7), redFluor 710-CD45R (B220) (RA3-6B2), Phycoerythrin (PE)-CD127 (IL-7Ra) (A7R34), violetFluor 450-CD44 (IM7), PE-Cy7-IFN-γ (XMG1.2), PE-Cy7-CD62L (L-Selectin) (MEL-14), and fluorescein isothiocyanate (FITC)-TCR β (H57-597) were from Tonbo Biosciences (San Diego, CA); PerCP-Cy5.5-KLRG1 (2F1/KLRG1), Alexa Fluor 700-TNF-α (MP6-XT22), APC-IL-17A (TC11-18H10.1), PE-IL-10 (JES5-16E3) were from BioLegend (San Diego, CA); eFluor 450-TCRγδ (eBioGL3 (GL-3, GL3)), was from eBiosciences (San Diego, CA). Cells were surface stained with the appropriate antibodies in phosphate-buffered saline (PBS), in the presence of Fc Block (BioLegend) and fixable viability dye (Ghost Violet 510) (Tonbo Biosciences). For intracellular cytokine staining (ICS), cells were stimulated as indicated, surface stained, then were fixed at room temperature with fixation buffer (BioLegend), and permeabilized with permeabilization buffer (BioLegend), and stained with the appropriate antibodies. Flow cytometry was performed with a BD LSRFortessa (BD Biosciences), and data were analyzed in FlowJo (Tree Star, Ashland, OR).

### Statistical analysis

For comparison of the means for two groups a Student’s t tests was performed using two-tailed analysis. For comparison of multiple groups, an ANOVA was initially performed and if significant differences among all the groups was noted, a Tukey’s test to adjust for multiple comparison was used. Data are presented as means and standard deviations. Incidence data were compared by Fishers’ exact test. A P value < 0.05 was considered significant.

### Data Availability

The authors declare that the data supporting the findings of this study are available within the paper.

## RESULTS

### Generation of VC2-derived influenza virus vaccine

HA protein is the major antigenic target of current influenza vaccines. To determine the potential of VC2 as a live vaccine vector against influenza virus infection we generated a recombinant VC2 virus which expresses influenza PR8 HA. Since glycoprotein C (gC) is a nonessential gene for HSV virus replication *in vitro*, we replaced gC with HA from the PR8 strain influenza virus (Fig. 1a). To enhance the expression of HA protein, we replaced the entire gC gene with PR8 HA and a pCAGGS promoter fused at its N terminus since pCAGGS promoter is a strong mammalian Pol II promoter. This virus was designated VC2-HA. We predicted that this promoter will drive higher HA expression compared to the original HSV gC promoter. The HA insertion was verified by PCR-assisted sequencing of the insertion site. VC2-HA was able to express PR8 HA in Vero cells, as indicated by the detection of HA with HA-specific antibody, PY102 (Fig. 1b). To ensure that insertion of HA did not affect virus replication we performed a multi-step growth curve to compare replication of VC2-HA to the parental VC2 virus in Vero cells at an MOI 0.01. Both VC2-HA and parental VC2 viruses exhibited a similar growth pattern (Fig. 1c).

**Figure 1.**
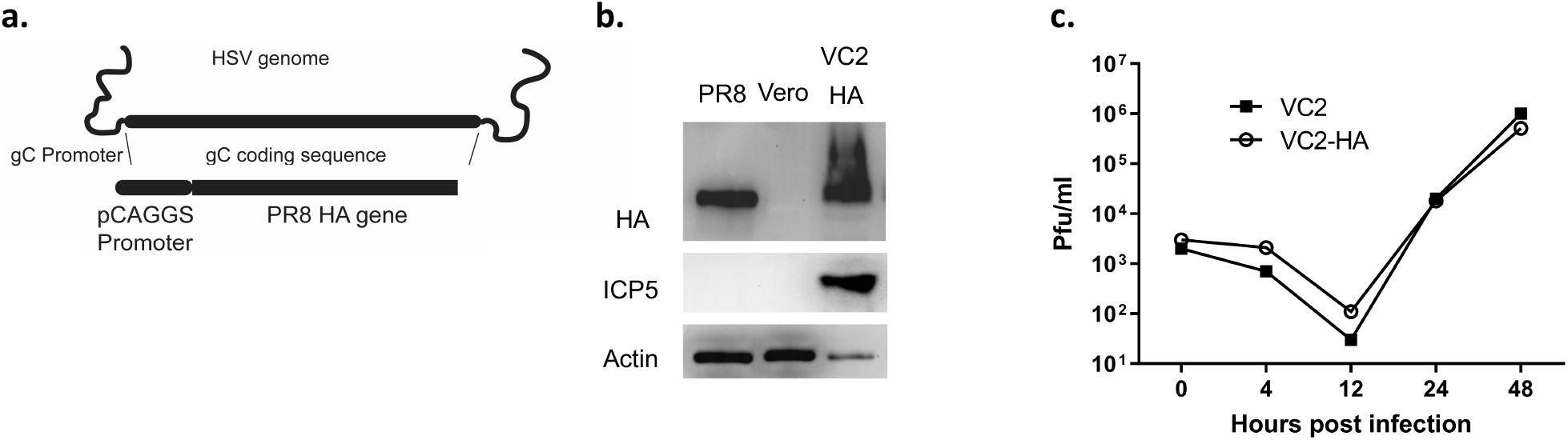
Construction of HSV-1-derived influenza vaccine. **a.** Construction of VC2-HA. **b.** Western blot showing expression of HA in VC2-HA infected Vero cells. Blot was stained with antibody to HA, viral capsid (ICP5), and actin. PR8 was used as a positive control and uninfected Vero cells were used as a negative control. **c.** Growth curve of VC2-HA and parental VC2 viruses. Vero cells were infected at a multiplicity of infection of .01. No significant differences in growth were observed.

### Immunization with VC2-pCA-HA protects mice from lethal challenge with PR8 virus

To evaluate the protection efficacy of recombinant VC2 expressing PR8 HA, we performed a lethal challenge mouse study using the VC2-HA vaccine. Specifically, 6-8-week-old Black 6/J mice (n=5) were vaccinated intramuscularly with VC2-HA at 1X10^5^, or 1X10^6^ PFU or VC2 parental virus at 1X10^6^ PFU per animal. Previous studies have shown that intramuscular route of inoculation with VC2 is safe in murine, guinea pig and non-human primate models and elicits immune responses to targeted antigens expressed in VC2 viruses (33, 40–42). For these experiments we used a single vaccination regimen. During the vaccination period there were no significant weight differences among the groups (data not shown). Three weeks after vaccination, mice were challenged intranasally with 1X10^3^ PFU of influenza A PR8 H1N1 virus. The two groups of mice vaccinated with different doses VC2-HA did not exhibit any weight loss or clinical signs, such as fur ruffling and heavy breathing during the two weeks post challenge (Figure 2b). However, the group of mice vaccinated with parental VC2 virus exhibited significant weight loss after the challenge and all eventually succumbed to infection (Figure 2b,c). The results indicated that vaccination of VC2-HA protected mice from the lethal challenge of PR8 viruses.

**Figure 2.**
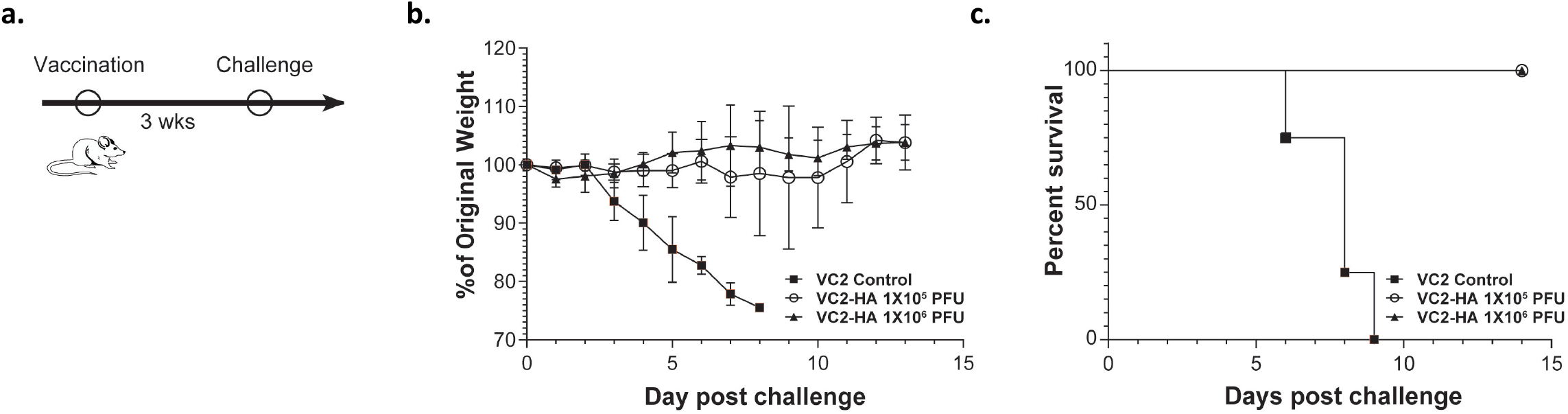
Vaccination of mice with VC2-HA protects mice from lethal challenge with influenza virus. **a.** Vaccination strategy. 6-8-week-old C57BL/6 J mice (n=5) were vaccinated intramuscularly with either 1X10^5^, or 1X10^6^ PFU VC2-HA. As a negative control, mice were vaccinated intramuscularly with 1X10^6^ PFU of VC2. For these experiments we used a single vaccination. Three weeks post vaccination mice were challenged intranasally with 1X10^3^ PFU of influenza A PR8 H1N1 virus. Mice were observed for weight loss (**b.**) and survival (**c.**). *p* < *0.05, **0.01, ***0.001, NS= not significant, by unpaired student’s *t* test.

### Vaccination of mice with VC2-pCA-HA induced PR8 HA specific antibodies with neutralizing capability

To evaluate the immunogenicity of recombinant VC2-HA, we first investigated the humoral responses induced by vaccination of mice with VC2-HA virus. 6-8-week Black/6J mice were vaccinated with VC2-HA at 1X10^5^ PFU, or VC2 parental virus at 1X10^6^ PFU per mouse intramuscularly since the intramuscular vaccination of VC2. Sera were collected from mice before prime, pre-boost (after prime) and three weeks after the boost for in vitro serological assays (Fig. 3a). Serum PR8 HA specific IgG titers were measured by ELISA using plates coated with purified PR8 viruses. The endpoint titers of serum IgG were used as the readout (Fig. 3a). After priming, all the VC2-HA vaccinations induced PR8 HA specific antibodies, whereas VC2 vector control group exhibit negligible PR8 HA binding signals.

**Figure 3.**
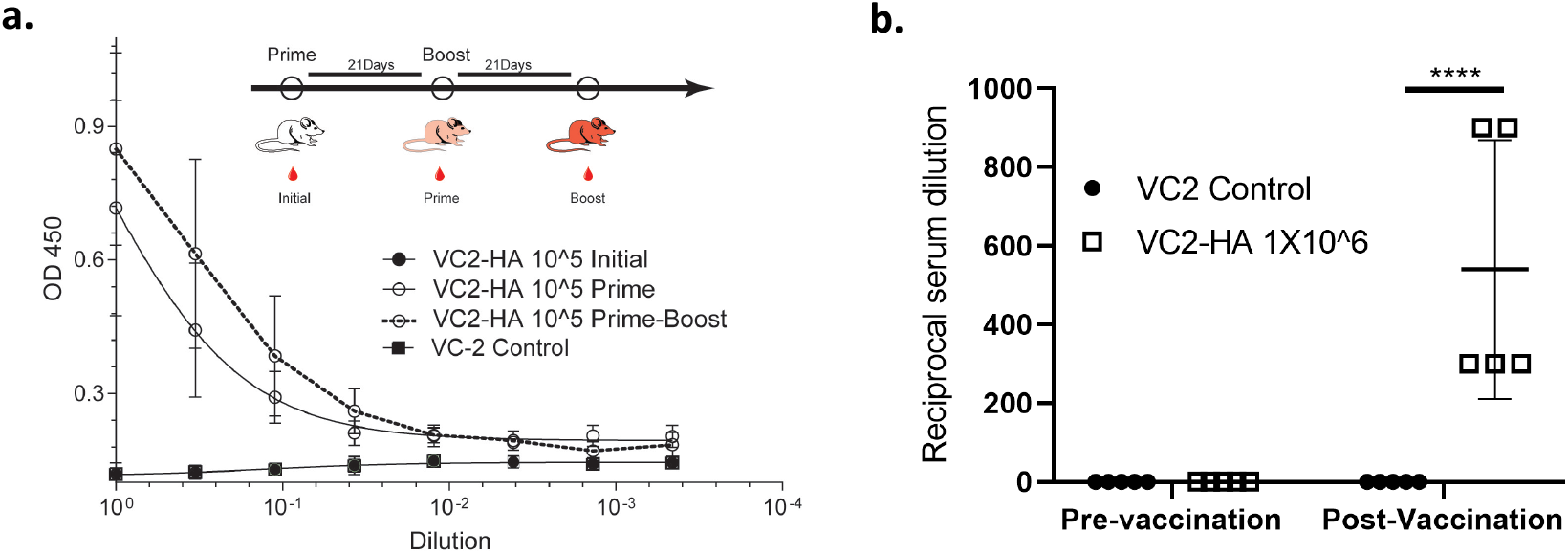
VC2-HA vaccination induces HA specific antibody responses. 6-8-week C57BL/6 J mice were vaccinated intramuscularly with VC2-HA at either 1X10^5^PFU or 1X10^6^ PFU, or VC2 vector alone at 1X10^6^ PFU. For these experiments we used a prime and a boost immunization after three weeks. Sera were collected from mice before prime, pre-boost (after prime) and three weeks after the boost. **a.** HA-specific IgG titers in sera were measured by ELISA using plates coated with purified PR8 viruses (Fig. 2a). **b.** Microneutralization Assays. *p* < ****0.0001 by unpaired student’s *t* test. N = 5. Data represent results of at least three independent experiments.

To evaluate the neutralizing activity of the antibodies induced by vaccination with VC2-HA, we performed microneutralization assays against influenza H1N1 PR8 virus since it carries the parental HA protein expressed in recombinant VC2 virus. For these experiments we used a single vaccination with 1X10^6^ PFU of VC2 HA or VC2. Only sera from animals vaccinated with VC2-HA exhibited neutralizing activity against PR8 virus (Fig. 3b). Collectively, the vaccination with VC2-HA elicited PR8 HA specific antibodies with robust neutralizing activity against PR8 virus.

### Vaccination of mice with VC2-pCA-HA induced PR8 HA specific T-cells

There is increasing evidence that the induction of T cells is important for the broadly protective properties of influenza vaccines (43). T cell responses to either HA or inactivated influenza virus were measured using splenocytes isolated 9 days post boost from either VC2 or VC2-HA vaccinated mice. Splenocytes were activated with either HSV-1 specific peptide as a control, or inactivated PR8 virus particles. gB_498-505_ is an immunodominant HSV-1 epitope for C57BL/6 mice (44). CD8+ T-cells from both VC2 and VC2-HA vaccinated mice were activated with gB_498-505_ (Fig. 4a). In contrast, only mice vaccinated with VC2-HA responded with IFN-γ and TNF-α expression after stimulation with heat-inactivated PR8 (HI PR8) virus (Fig. 4a) and expanded upon re-stimulation by HA protein (Fig. 4b). Furthermore, we observed an increase in CD4+ and CD8+ T cells with an effector/memory phenotype (CD44high CD62L-) in VC2-HA vaccinated mice, as compared to VC2 vaccinated controls (Fig. 4b).

**Figure 4.**
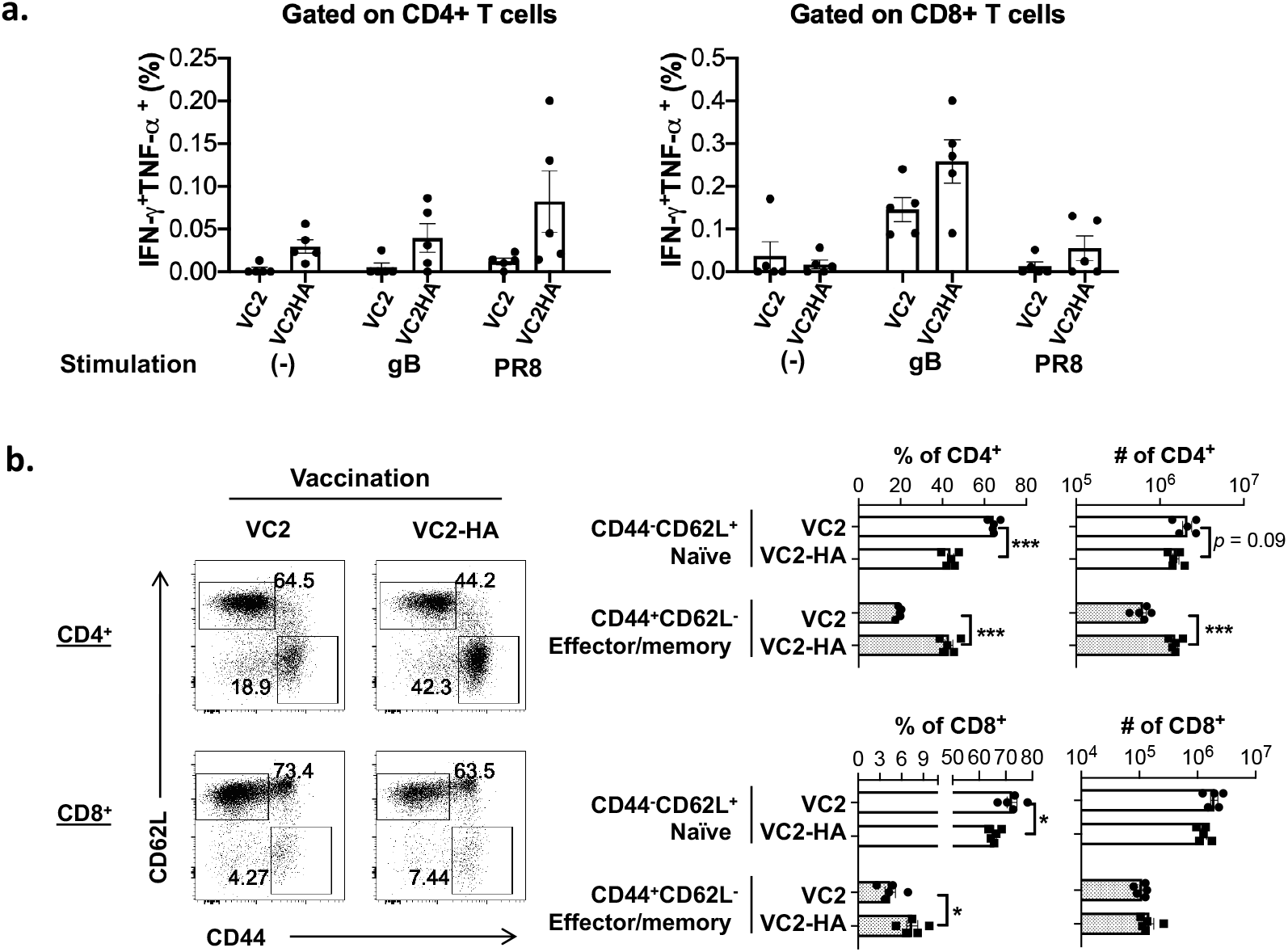
VC2-HA vaccination induces both CD4+ and CD8+ effector/memory T-cells. Splenocytes from vaccinated mice were analyzed 9 days post boost. **a.** Percentages of IFNγ and/or TNFα-producing CD4+ and CD8+ T cells post re-stimulation with HSV-1 gB_498-505_ peptide or heat-inactivated PR8 virus for 5 hours in vitro. **b**. Representative FACS plots and summary of percentages of naïve and effector/memory CD4+ and CD8+ T cells. Summary of percentages of expanded CD4+ and CD8+ T cells post re-stimulation. *p* < *0.05, **0.01, ***0.001, NS = not significant, by unpaired student’s *t* test. N = 5. Data represent results of at least three independent experiments.

## Discussion

Previous studies from several groups have established that HSV-1 is a safe and effective live vaccine vector, which triggers robust immune responses against multiple infectious pathogens including HIV and rotavirus, among others (45–48). Herein, we generated a HSV-1-derived vaccine against influenza virus, a significant human and animal respiratory viral pathogen. We engineered the recombinant HSV-1 VC2 strain to express the influenza HA protein under a strong promoter. Immunization of mice with this novel vaccine induced anti-influenza immune responses and that a single vaccination with VC2 HA protects mice against lethal influenza virus challenge.

The utility of viruses as vectors for vaccines requires that they are safe, immunogenic, and capable of inducing protection in the presence of pre-existing immunity. Our novel vector, VC2 possesses mutations that abrogate infection of neurons via neuronal axons (34). This is a rational design that disrupts the ability of the virus to establish latency. The inability to establish latency is expected to preclude the majority of clinical symptoms associated with HSV-1 infection. VC2 has been shown to be safe and immunogenic in murine (including SCID mice), guinea pig and non-human primate models (33, 38–41). Our data here show that VC2-HA promotes strong humoral and cell mediated immune responses and protects mice from lethal influenza virus challenge. Consistent with this, we have previously shown that VC2 can be used to create a safe, immunogenic vaccine against equine herpesvirus 1 (42).

All viral vector-mediated therapeutic or intervention strategies can be potentially affected by pre-existing immunity in the target population. This is a concern for HSV-1-derived vectors due to their high prevalence in human populations. However, it is known that individuals are capable of being re-infected by differing strains of HSV throughout their lifetime (48). Further, a hallmark of herpesvirus infection is their ability to reactivate and spread in the presence of sizable host anti-herpesvirus immune responses (49). These characteristics of the natural history of herpesvirus infection, as well as a number of studies demonstrating that pre-existing immunity had no substantial effect on vector efficacy (38–42) inform the usage of herpesvirus-derived vectors for use in anti-cancer and vaccine applications. Specifically, clinical trial data from T-Vec^™^, the FDA approved HSV-1-derived oncolytic virotherapy, found no difference in treatment efficacy between seropositive and seronegative individuals. Consistent with this, we and others have found no effect of HSV-1 seropositivity on the efficacy of oncolytic virotherapy in mice (29, 49, 50). While the current study was not performed on HSV-1 seropositive mice our data shows we are able to get increased anti-HA immune responses after a prime and boost compared to a single vaccination. As we were unable to detect HA on virion particles of VC2-HA (data not shown), this suggests VC2-HA boost virus was able to infect and express HA in seropositive mice.

There are a number of groups currently working on HSV-vectored vaccines against a number of human and animal pathogens. Multiple approaches ranging from HSV based amplicons, replication defective, attenuated or conditionally defective vectors have been considered (45–48). Our approach to altering the pathogenic potential of HSV-1 is unique in that it allows the virus to replicate essentially as a wild type virus without the potential for causing disease as seen in other, even attenuated viral vectors. This wild type like replication is important as it likely allows for the maximum potential immunogenicity of the vectored antigen. Furthermore, we have shown that VC2 possesses the gK31-68 mutation that forces the virus to enter only via endocytosis instead of fusion of the viral envelope with cellular plasma membranes (51). This altered mode of entry prevents the virus from entering into neuronal axons as well as potentially altering innate immune responses and downstream adoptive immune responses as evidenced by the fact the the VC2 intramuscular immunization was more effective in conferring against lethal ocular challenge with the human clinical strain HSV-1 (McKrae) (39). In this regard, VC2 appears to possess an “adjuvant” effect that can facilitate robust immune responses against heterologous antigens expressed via the VC2 vector.

Current influenza vaccines induce neutralizing antibodies as their primary mechanism of protection. These vaccines induce strain-specific immunity and as a result leave human populations at risk for infection with other influenza strains. It has been suggested that a T-cell response may be necessary to induce broad protection against multiple strains of influenza (9). Our data demonstrate that our vaccine is capable of producing both cellular and humoral anti-influenza immune responses. With our approach, in addition to neutralizing antibodies, we achieved significant induction of anti-influenza CD4+ and CD8+ T-cells. In this study, we saw greater induction of anti-influenza CD4+ T-cells than CD8+ T-cells. We believe that this is likely due to the use of inactivated virus and recombinant HA protein to stimulate splenocytes in our experiments. It is possible that HA peptide libraries would be more effective at identifying HA specific CD8+ immune responses. Regardless, our data demonstrate that VC2-HA is capable of inducing significant influenza specific T-cell responses. In future experiments it will be critical to test the ability of our vaccine to induce broad protection against multiple strains of influenza.

Herein, we present data that the novel HSV-1-derived vaccine vector VC2 expressing the influenza virus HA antigen induces a robust influenza-specific humoral and cellular immune responses, resulting in protection of mice from lethal influenza virus challenge. These observations support continued development of VC2-HA to better characterize the breadth of protection elicited by this vaccine against heterotypic strains of influenza. Additionally, these data support the utility of the VC2 vectored vaccine approach for the further development of a universal influenza vaccine as well as vaccines against significant emerging and re-emerging pathogens.

## Acknowledgements

This research was supported in part by NIH 1R21AI147057 grant to R.H. and a grant from Louisiana Board of Regents Governor’s Biotechnology to K.G.K and Core Facilities supported by NIH:GM103424 and NIH GM110760. I.K.U. is supported by a graduate stipend from LSU School of Veterinary Medicine.

## Contributions

The overall study design was by R.H., P.J.F.R., and K.G.K. Experiments and analyses were performed by P.J.F.R., H.D., I.K.U., M.C.M., B.B., G.S.T., W.H. and R.H. Animal Studies were performed by R.H., P.J.F.R., H.D., I.K.U., and M.C.M. The manuscript was written by R.H. and P.J.F.R., with input and comments from all authors.

